# Isotope effects on radical pair performance in cryptochrome: a new hypothesis for the evolution of animal migration

**DOI:** 10.1101/2022.08.08.503164

**Authors:** Ismael Galván, Abbas Hassasfar, Betony Adams, Francesco Petruccione

## Abstract

Mechanisms occurring at the atomic level are now known to drive processes essential for life, as revealed by quantum effects on biochemical reactions. Some macroscopic characteristics of organisms may thus show an atomic imprint, which may be transferred across organisms and affect their evolution. This possibility is considered here for the first time, with the aim of elucidating the appearance of an animal innovation with an unclear evolutionary origin: migratory behaviour. This trait may be mediated by a radical pair (RP) mechanism in the retinal flavoprotein cryptochrome, providing essential magnetic orientation for migration. Isotopes may affect the performance of quantum processes through their nuclear spin. Here, we consider a simple model and then apply the standard open quantum system approach to the spin dynamics of cryptochrome RP. We changed the spin quantum number (*I*) and g-factor of hydrogen and nitrogen isotopes to investigate their effect on RP’s yield and magnetic sensitivity. Strong differences arose between isotopes with *I*=1 and *I*=1/2 in their contribution to cryptochrome magnetic sensitivity, particularly regarding Earth’s magnetic field strengths (25-65 µT). In most cases, isotopic substitution improved RP’s magnetic sensitivity. Migratory behaviour may thus have been favoured in animals with certain isotopic compositions of cryptochrome.

## 1. Introduction

Life depends upon interactions between the constituent molecules of organisms. Isolated molecules are inanimate. But the physicochemical characteristics of biomolecules, which determine their interactions, are given by their constituent atoms [1]. Life therefore occurs, to some extent, at an atomic scale. Not all atoms of a given element are equal: elements occur as isotopes, i.e., atoms of the same element that differ in the number of neutrons in their nuclei, thus having different mass numbers and nuclear spins. The isotopic composition of molecules might be transferred across molecules, and across organisms, during molecular biosynthesis independently of the genetic code, as nucleic acids cannot determine their own isotopic composition [2]. Atomic properties like this may be responsible for some macroscopic characteristics of organisms, but this possibility has been only envisaged by some authors [3-5]. However, if the isotopic composition of biomolecules is responsible for macroscopic characteristics of organisms, this may have profound consequences for organic evolution.

Different isotopic compositions of the same molecule (i.e., isotopologues) do not behave equally in some chemical reactions [6]. Stable isotopes, i.e., those not undergoing radioactive decay, are particularly relevant for organisms, because the nuclei of the most common biological elements (C, H, O, N and S) exist as 13 stable isotopes, making those biomolecules appear as mixed isotopologues. The different mass numbers of these stable isotopes give them slightly different properties, leading to their partial separation during chemical, biological and physical transformations in what is termed isotope fractionation. Fractionation occurs because of the influence in chemical reactions of the different masses (nuclei with more neutrons are heavier), volumes (nuclei with more neutrons are bigger) and spins (nuclei with certain spin values have a magnetic moment) of isotopes [7]. This explains the distribution of isotopes in the biosphere, as fractionation occurs first in the environment, then during plant photosynthesis, and lastly during animal metabolic reactions [8].

Different experiments have proved that organisms grown under certain isotopic media or given diets with certain isotopes experience oxidative stress and resistance to change the isotopic composition of their tissues [9]. This has been observed at all levels of complexity, from microorganisms to vertebrates such as birds and humans, and in isotopes of elements such as H, C and N [10-12]. These studies indicate that organisms have a preferred isotopic environment, because there is an optimum isotopic composition of molecules under which biochemical reactions accelerate [9,13,14]. Concomitantly, an evolutionary optimization explains the negative physiological effects that organisms experience under deviations of their isotopic composition, even if adaptation can be achieved [9]. Chemical and evolutionary optimizations thus make likely that every organism has a specific isotopic profile that is not randomly determined by environmental influences.

Yet, a small proportion of variation in the isotopic composition of animals is due to the incorporation of isotopes with the food [15,16]. Ecological studies even assume that the whole-body composition of an animal closely resembles that of its diet when tracking the trophic behavior of animal groups within a given species [8]. This is used to infer the trophic niche (isotopic niche or “isospace”) of species of animals by plotting the composition of certain tissues in C isotopes against that of N isotopes. Isospaces show significantly differentiated, characteristic isotopic compositions of many species of animals, including hominids [17,18]. Importantly, isospaces do not overlap even between coexisting species that are morphologically and ecologically similar [19]. This supports the idea that, even considering the dietary source of variability in the isotopic composition of animals, every species may have a characteristic isotopic composition. This actually agrees with the existence of niche conservatism, i.e. the retention of ecological traits in species across space and time, as revealed by non-overlapping isospaces in ecologically similar species of animals [19,20]. These differentiated isospaces of animals have never been considered an intrinsic trait of species, but here it is hypothesized that the isotopic composition of molecules might be more similar within animals belonging to the same species than between animals of different species (i.e., interspecific variability > intraspecific variability). Indeed, even if part of variation in the isotopic composition of animals is due to dietary incorporations, individuals of the same species share biochemical reactions of metabolism, thus experiencing isotope fractionation processes that must be more similar than in individuals of other species, which possess different metabolic profiles. This is because metabolic pathways are shared by all individuals of the same species, an observation that has allowed the development of metabolomics, in a similar way as genomics, as a useful tool to elucidate the evolution of species-specific traits in organisms [21,22]. Certain isotopic compositions in certain species may therefore favour some biochemical reactions and thus promote the evolution of important phenotypic traits.

The development of the field of quantum biology suggests that biochemical reactions involved in physiological processes are mediated by quantum mechanisms [23,24]. One such mechanism is represented by radical pairs (RP), which are strongly influenced by isotopes [25] and are potentially responsible for animal magnetoreception: the ability to perceive the Earth’s magnetic field and use it to orientate during migratory movements. This ability is most notably observed in birds, but has evolved in all major vertebrate groups and some invertebrates [26,27]. The mechanism of magnetoreception and the ability for magnetic orientation was proposed to be mediated by a light-initiated RP in the eyes of birds, whose recombination disruption by the magnetic field is recognized by the bird’s nervous system [28]. This is currently the leading theory of avian magnetoreception. Cryptochrome 4 (CRY4), a pigment belonging to the only vertebrate protein class that forms radical pairs upon photo-excitation (flavoproteins), has been suggested to be the likely magnetoreceptive protein, localized in double-cones and long-wavelength single cones in the retina of birds [29]. Recent research has demonstrated that the photochemistry of CRY4 in night-migratory birds is more magnetically sensitive than that of some non-migratory species [30]. However, the exact type of cryptochrome responsible for magnetoreception remains a subject of debate. There are five different types of cryptochrome found in birds’ retinas. In addition to CRY4, cryptochrome 1a (CRY1a) has been put forward as the likely site for a light-sensitive magnetic compass, due to its specific location in the outer segments of the ultraviolet cone cells of birds [31]. The majority of research into the role of RPs in magnetoreception has focused on birds. The RP mechanism could, however, be the same in other animals with magnetic orientation capabilities, as the expression of magneto-sensitive cryptochrome has been shown to occur in the retina of insects [32], and even of humans [33]. The capacity for magnetic orientation has also been shown to reside in the eyes of other mammals [34].

In its simplest iteration, an RP consists of two radicals (molecules with an odd number of electrons) that have been created simultaneously by chemical bond breaking or light-induced electron transfer, forming a spatially separated but spin-correlated and possibly entangled electron pair. This entanglement is due to the fact that the RP is often assumed to be created in a singlet, or maximally entangled state, although other options have been discussed [35]. Due to external and local (hyperfine interaction) magnetic fields, the spin character of the radical pair is coherently interconverted between singlet (anti-parallel electron spins) and triplet (parallel electron spins) states [36]. The spin state of the electron (as a purely quantum mechanical concept) then influences subsequent chemical reactions, converting information about the magnetic environment into a chemical signature. As outlined previously, this mechanism has been hypothesized to be responsible for magnetoreception in birds: cryptochrome in the retina is photoexcited by light, forming an RP that is initially in the singlet state and is interconverted between singlet and triplet states by hyperfine coupling. The rate of recombination of singlet states also depends on the orientation of the Earth’s magnetic field, and the interconversion is oscillatory coherent, generating a quantum beat that is used by birds for orientation [37]. See Figure 1 for details of the RP formation.

**Figure 1.**
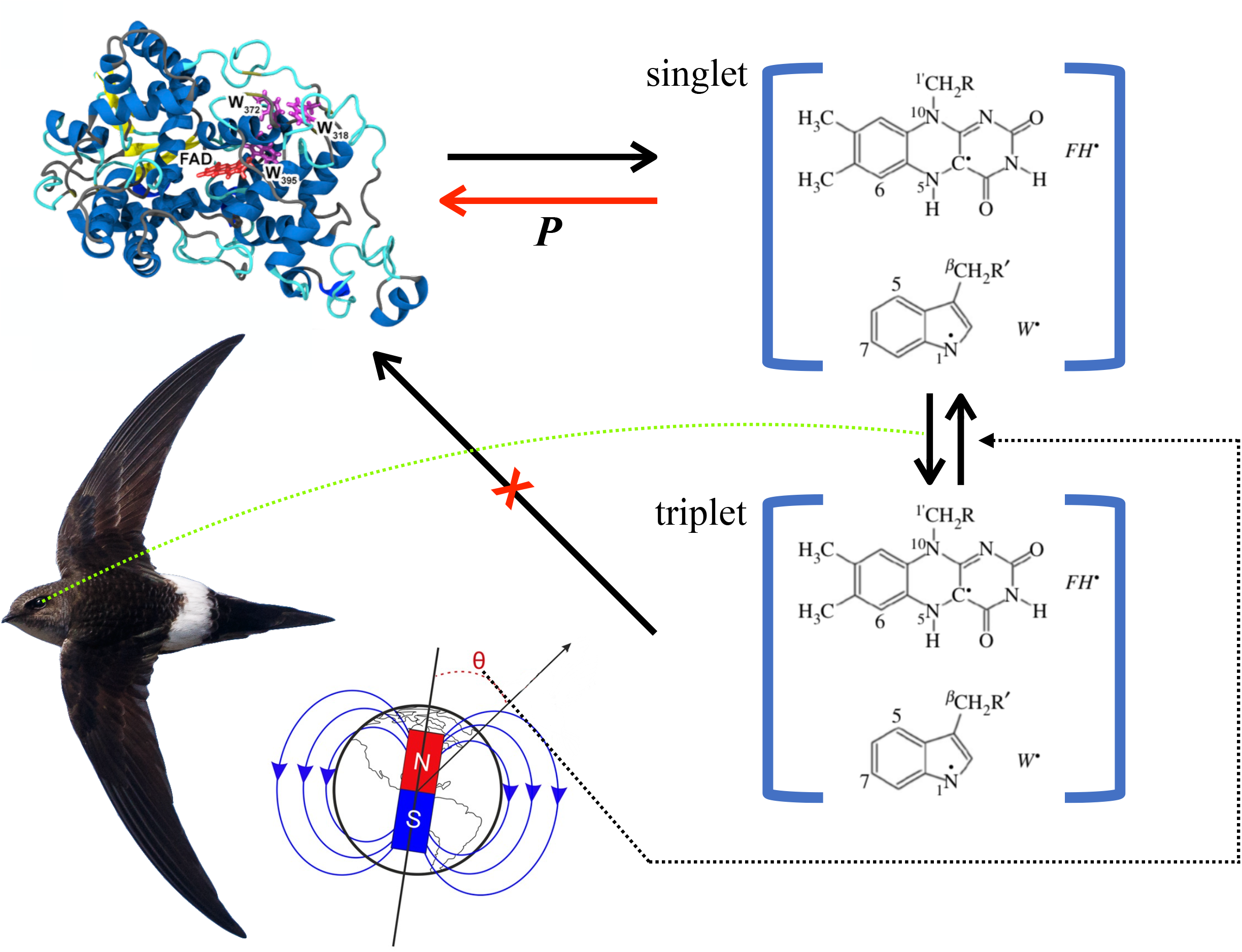
The radical pair (RP) mechanism in cryptochrome may facilitate magnetoreception in migratory animals. Within the protein cryptochome 4, a flavin residue and a tryptophan form a dyad that, upon photoexcitation, gives rise to a flavin radical (*FH*^•^, or FAD^•−^) and a tryptophan radical (*W*^•^, or TrpH^•**+**^). Then the RP can either produce photoproducts, or recombine to produce the original, zero-spin dyad molecule. The radical pair is initially in singlet state, but hyperfine coupling leads to conversion into non-reactive triplet states. The recombination reaction thus requires a triplet-singlet spin conversion, which is a function of the magnetic field, the nuclear spin and its projection, the nuclear magnetic moment, the hyperfine coupling constant, the frequency and amplitude of the magnetic field, and the interaction between unpaired electrons [27]. As the nuclear spin differs between isotopes, isotopes with specific *I* values experience specific hyperfine coupling between the unpaired electron and the magnetic nucleus, potentially leading to fast triplet-singlet conversion. This conversion is slower in isotopes with other *I* value. In the flavin-tryptophan radical pair, the triplet-singlet conversion rate depends on the orientation of the Earth’s magnetic field, allowing orientation and migratory movements. Molecular and Earth images taken from Dreyer et al. [27] (open access content). Swift image belongs to Stuart Price (https://flic.kr/p/TLn66s) and is under CC BY-SA 2.0 license (https://creativecommons.org/licenses/by-sa/2.0/).

Magnetoreception in birds is strongly believed to be an RP mechanism based on retinal cryptochrome [38]. However, the strong influence that isotopes exert on RP mechanisms has been largely overlooked, with the exception of Player and Hore [39]. This influence is due to the fact that one of the dominant interactions involved in RP dynamics is the internal magnetic interactions, known as hyperfine interactions, which are caused by the surrounding nuclear spins that can interact with an unpaired electron. Nuclei can have no spin (e.g., ^12^C and ^16^O), integer spin (e.g., ^14^N and ^2^H) and also half-integer spin (e.g., ^13^C and ^1^H) [40]. Thus, different isotopes of an element can have different magnetic effects on the dynamics of the system. An RP with isotopes with certain spin undergoes fast triplet-singlet conversion, while this is strongly delayed in the case of isotopes with other spins [25]. Cryptochrome molecules with isotopes with certain spin numbers might thus impair the RP mechanism, which might be favoured in cryptochrome isotopologues with other spin numbers (Figure 1). The ability for magnetic orientation and long-distance migration may therefore be more likely to arise in animals that have inherited cryptochrome molecules with higher proportions of isotopes with certain spin numbers.

Here we hypothesize that the ability to conduct long-distance migration has evolved only in animals with a prevailing contribution of isotopes with certain spin to the cryptochrome molecule. To evaluate the potential of this hypothesis to explain interspecific variability in migratory ability, we modeled spin dynamics of the broadly accepted RP model of the cryptochrome-based avian compass.

## 2. Methods

We considered two different theoretical scenarios to investigate our hypothesis. Both involve the conventional radical pair approach, though there is some suggestion that adding a third radical improves the performance of the radical-based compass [41]. In the first scenario (**Scenario 1**), we explored a simple model with one hyperfine-coupled nucleus for the calculation of the magnetic sensitivity (MS) of cryptochrome depending on different quantum spin numbers associated to different nuclei. We explored MS in a different range of anisotropic hyperfine coupling constants in the presence of an external magnetic field, similar to the Earth’s geomagnetic field. The second scenario (**Scenario 2**) looks more closely at isotope substitution of the specific model by changing the spin quantum number and also the g-factor for scaling the strength of hyperfine interaction for different geomagnetic field intensities. In this second scenario, we used the DFT calculations of nuclear hyperfine coupling tensors by Hiscock et al. [42] for cryptochrome, in addition to a simpler version with up to three nuclei that has previously been used by Jain et al. [43] and that assumes that all hyperfine tensors are simultaneously diagonalized. In this second scenario, we show that both singlet yield and MS change with isotope substitution.

The spin dynamics of RPs are described by the following Hamiltonian:

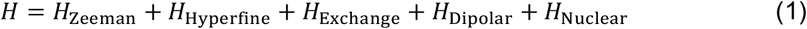

Among these five terms, the exchange, dipolar and nuclear Zeeman interactions usually are negligible as compared to the other two terms [44].

Two dominant interactions are conventionally taken to be involved in RP dynamics: one of them is the Zeeman interaction (*H*_Zeeman_), due to the magnetic field of an external source (e.g., the geomagnetic field). The other is due to internal magnetic interactions, known as hyperfine interactions (*H*_Hyperfine_) which are caused by the local surrounding nuclear spins that can interact with an unpaired electron. The RP Hamiltonian with one nucleus can thus be written as:

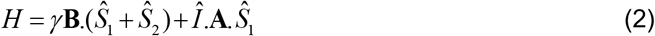

where ‘I’ is the nuclear spin operator, 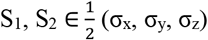 are electron spin operators (σ_x_, σ_y_, σ_z_ are the Pauli matrices), γ = µ_0_g is the gyromagnetic ratio, µ_0_ is the Bohr magneton, and g is the electron g factor (= 2.0023). We will consider the simplest case in which the hyperfine interaction is anisotropic, A = diag(0, 0, a) as well as the more complex case of A = diag(a_x_, a_y_, a_z_). The external field (geomagnetic field) is characterized by B = B_0_(sinθcosφ, sinθsinφ, cosθ); B_0_ = 47 µT is the local geomagnetic field, and θ is the magnetic field orientation with respect to the molecular axis [45]. These interactions dictate the dynamics of the electrons’ spin state and can cause coherent conversion between singlet and triplet states.

Different approaches have previously been used to tackle RP dynamics and recombination. These include Haberkorn’s [46] master equation, the Jones-Hore measurement master equation [47], Kominis’ [48] measurement master equation approach, which commonly includes the Liouville-von Neumann equation, as well as other open quantum systems approaches [49]. There remains some debate as to which is the best approach [44]. We use the Markovian quantum master equation approach to simulate the dynamics of the RP system of cryptochrome, similar to Gauger et al. [50]. In this quantum master equation approach, which conserves probability unlike the Liouville equations, we use the ‘shelving states’ |*S* ⟩ and | *T* ⟩ to represent the singlet and triplet products that are spin-selected from the initial electronic singlet or triplet states of the RP. Mathematically, we use the direct sum to extend the Hilbert space of the radical pair to include the shelving states | *S* ⟩ and | *T* ⟩ as extra basis vectors. The recombination of the RP into singlet and triplet channels is modeled through decay operators. For example, for one nucleus model with I=1/2, there are eight projections, four projections for the ‘up’ (|↑>) state of nucleus 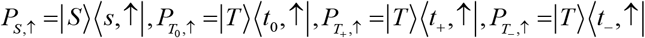, and, similarly, four projections for the ‘down’ (|↓>) nuclear state 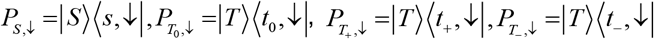 [49]. The singlet yield is defined as the proportion of recombined chemical products originating from the singlet precursor. One of the two events will occur, and the final populations of | *S* ⟩ and | *T* ⟩ give the singlet and triplet yield.

The standard, GKSL (Gorini–Kossakowski–Sudarshan–Lindblad equation) master equation can be defined as [51]:

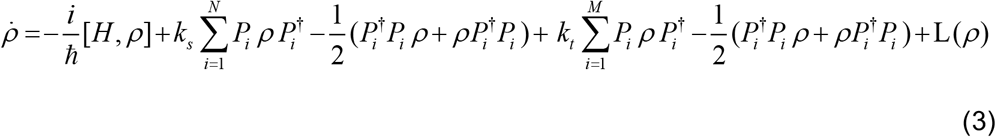

Where *P*_*i*_ are the projections on singlet and triplet, and N and M are the number of projection operators into singlet and triplet that depend on the Hilbert space of the nucleus. In the case of one nucleus with *I*=1/2, then N=2 and M=6, but in the case of one nucleus with I=1, then N=3 and M=9. L(ρ) is the standard Lindblad dissipator, used here to describe noise in the system. Several different noise models have previously been proposed, with different noise rates [50,52]. Here, we do not aim to explore all of the different noise models. As an important example we consider a special type of noise, perfectly correlated pure local dephasing noise, given by:

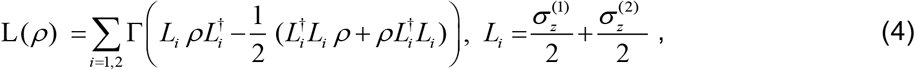

where Γ is the dephasing rate. Results suggest that with this noise model MS is quite robust and is even enhanced by the presence of correlated dephasing Cai et al. [52].

The master equation is given in terms of the rates of recombination to singlet and triplet products, k_s_ and k_t_, where H is given in Eq. (3). The dynamics of the master equation of the RP system were calculated using the QuTiP quantum toolbox in the Python module [53], which is developed for simulating quantum systems, particularly open quantum systems.

Here, we aim at showing that isotopes react preferentially (in terms of singlet or triplet yield and MS) depending on their nuclear spin quantum number I and strength of hyperfine interaction, which is embedded in their g-factor. We thus explored the quantum spin dynamics of the RP to investigate isotope-dependent magnetic field effects in the specific system of cryptochrome.

We only consider the spin dynamics, neglecting any effects related to nuclear mass and volume. This is possible due to the fact that, within the Born-Oppenheimer approximation, electronic properties (such as spin density) do not depend on the nuclear mass and nuclei are considered point charges in quantum chemistry, meaning that nuclear volume is also irrelevant [54]. Therefore, we need only to change the spin quantum number and the g-factor to explore the effect of isotope substitution (we explain in section 2.1 how g-factor affects hyperfine coupling constant).

1. We made the following assumptions for the sake of simplification: All projectors have the same recombination rates. We have used two different recombination rates: k_s_=k_t_=k=0.1 µs^-1^ and 0.5 µs^-1^ (τ = 1/k = 10 µs and 2 µs, the lifetime of the radical pair, τ, is defined as the reciprocal of k) [50]. It is clear from the results that the sensitivity of the avian compass relies on the recombination rate, where a longer lifetime allows for greater singlet-triplet interconversion and resultant greater sensitivity.
2. One electron of the pair has no hyperfine interaction and the other electron experiences a hyperfine interaction due to the surrounding nucleus.
3. The initial state of our model ρ_0_ assigns a pure singlet state 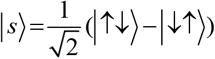 to the electrons, and a completely mixed state to the nucleus due to its interaction with the neighboring soft matter environment, initial state 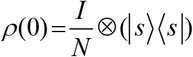 where N is the dimension of the Hilbert space of the nucleus (e.g., for one nucleus with spin 1/2, N=2, one nucleus with spin 1, N=3, and so on).
4. Without loss of generality, the axial symmetry of the hyperfine tensor allows us to assume that φ = 0 [50].

Our analyses are consistent with previous analyses of singlet yield for various hyperfine interaction strengths for different spin quantum numbers, and reproduce the sensitivity behaviour shown by Cai et al. [52] and Lee et al. [55] for cryptochrome. The magnetic field effect was calculated as Φ_S_, i.e., the fractional yield of the singlet reaction product once all radical pairs have reacted. Φ_S_ lies in the range [0,1] and is related to the triplet product yield by Φ_T_ = 1-Φ_S_. To quantify the effectiveness of a radical pair as a magnetic compass, we define the anisotropy of the reaction yield, i.e. MS, as 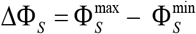 (the difference between the maximum and the minimum singlet yield as a function of the inclination). S denotes the singlet yield and the max/min are with respect to the magnetic field orientation, the difference between the maximum and minimum values of Φ_S_ being calculated as a function of the direction of the magnetic field vector, following the example of Lee et al. [55].

### 2.1. Isotope substitution

Substitution of one nucleus X, with spin quantum number I and factor g_n_, by its isotope X’ with I’ and g_n_’, changes the number of lines detected by Electron Spin Resonance Spectroscopy from (2I+1) to (2I’+1) and the coupling constant from a_X_ to a_X’_ = a_X_. g_n_’/ g_n_ [56]. Thus, for example, replacing a proton (*X* = ^1^*H* = *H* ; *I* = 1/ 2; *g*_*n*_ = 5.5854) by a deuteron (*X* = ^2^*H* = *D* ; *I* = 1; *g*_*n*_ ‘ = 0.8574) increases the number of lines from 2*1/2 + 1 = 2 to 2*1 + 1 = 3 and decreases the coupling constant from a_H_ to a_D_ = a_H_ (0.8574/5.5854) = 0.1535 a_H_. On the other hand, substituting a ^14^N nucleus (^14^*N* = *N* = *H*; *I* = 1; *g*_*n*_ = 0.4038) by its ^15^N isotope (^15^*N* = *N*’; *I* = 1/2; *g*’_n_ = -0.5664) decreases the number of lines from 2*1 + 1 = 3 to 2*1/2 + 1 = 2 and converts the coupling constant a_N_ into a_N_’ = a_N_ (−0.5664)/0.4038 = -1.4027 a_N_ [56] (Figure 2). The properties of H and N and their isotopes are summarized in Table 1. As previously stated, different isotopes of a given element can have different magnetic effects on the dynamics of the system. Apart from the spin quantum number, the strength of hyperfine interaction can be determined via the nuclear g-factor. By varying the number of nuclear spins interacting with the RP, it is possible to investigate the effects of different nuclear environments and thus draw some conclusions about the structure of cryptochrome, the biomolecule in which it is thought that the RP reaction takes place.

**Figure 2.**
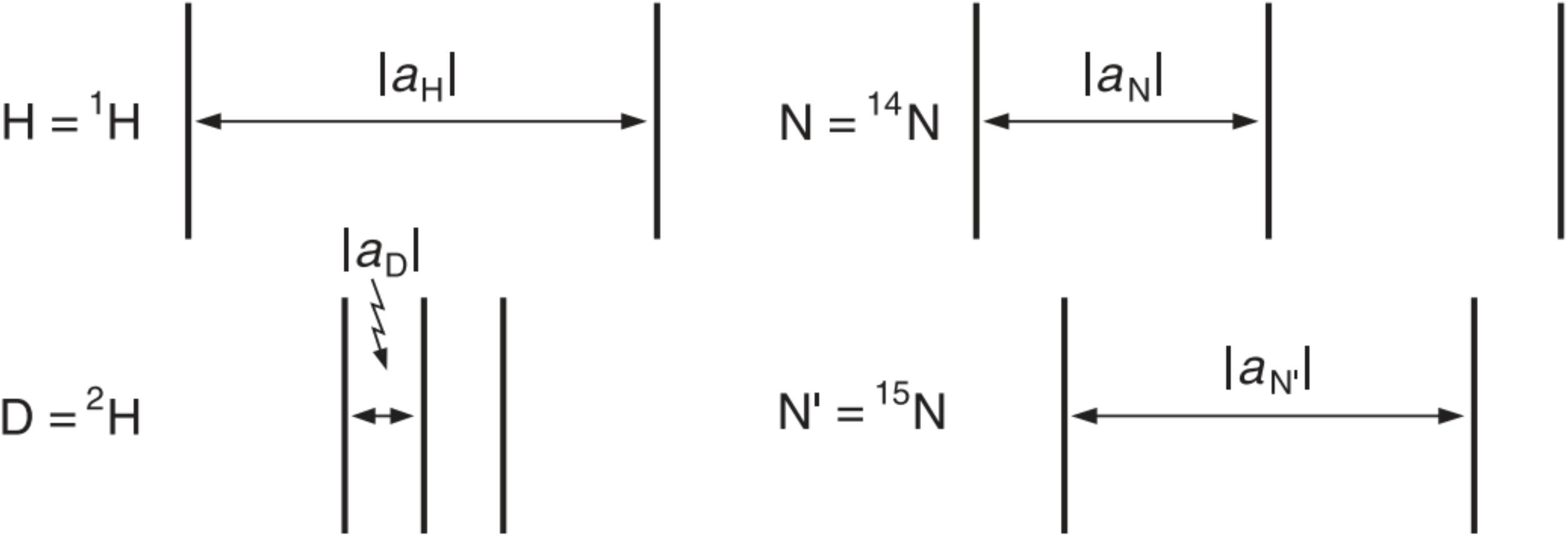
Hyperfine patterns occurring during the replacement of a proton (H) by a deuterium (D) and a ^14^N nucleus (N) by its ^15^N isotope. Following Gerson and Huber [56].

**Table 1.** Nuclear isotope effect of nitrogen and hydrogen.

| # | Name | Symbol | Z | N | A | Spin I<br>parity | ($\mu$ in $\mu_N$) | Nuclear<br>g factor<br>($g_N$) | Natural<br>abundance | Isotopic<br>mass ($u$) |
| --- | --- | --- | --- | --- | --- | --- | --- | --- | --- | --- |
| 1 | Hydrogen | H | 1 | 0 | 1 | 1/2 + | +2.7928 | +5.5857 | 99.9885 % | 1.0078 |
| 2 | Hydrogen | H (D) | 1 | 1 | 2 | 1 + | 0.8574 | +0.8574 | 0.0115 % | 2.0141 |
| 3 | Nitrogen | N | 7 | 7 | 14 | 1 + | 0.4037 | +0.4038 | 99.636 % | 14.0030 |
| 4 | Nitrogen | N | 7 | 8 | 15 | 1/2 - | -0.2832 | -0.5663 | 0.364 % | 15.0001 |

$\mu$ in $\mu_N$ : Nuclear magnetic moment of spin I in $h/2\pi$ .

## 3. Results

### 3.1. Scenario 1

In this scenario, the only difference between the treatment of nuclei in the model lies in their spin quantum numbers, meaning that we investigate MS with respect to the hyperfine coupling of a single nucleus model for two different spin quantum numbers (*I*=1 and *I*=1/2). We also consider only the simplest type of anisotropy, with A = (0, 0, a). In this simplest case, we do not include noise in the model. The results of calculations of MS for different strengths of anisotropic hyperfine coupling are shown in Figure 3 and show a marked advantage for spin *I*=1/2 than *I*=1. These results are also useful in that they give us an idea of the best and worst hyperfine coupling strengths with respect to magnetic sensitivity, albeit in the limited case of simple anisotropy.

**Figure 3.**
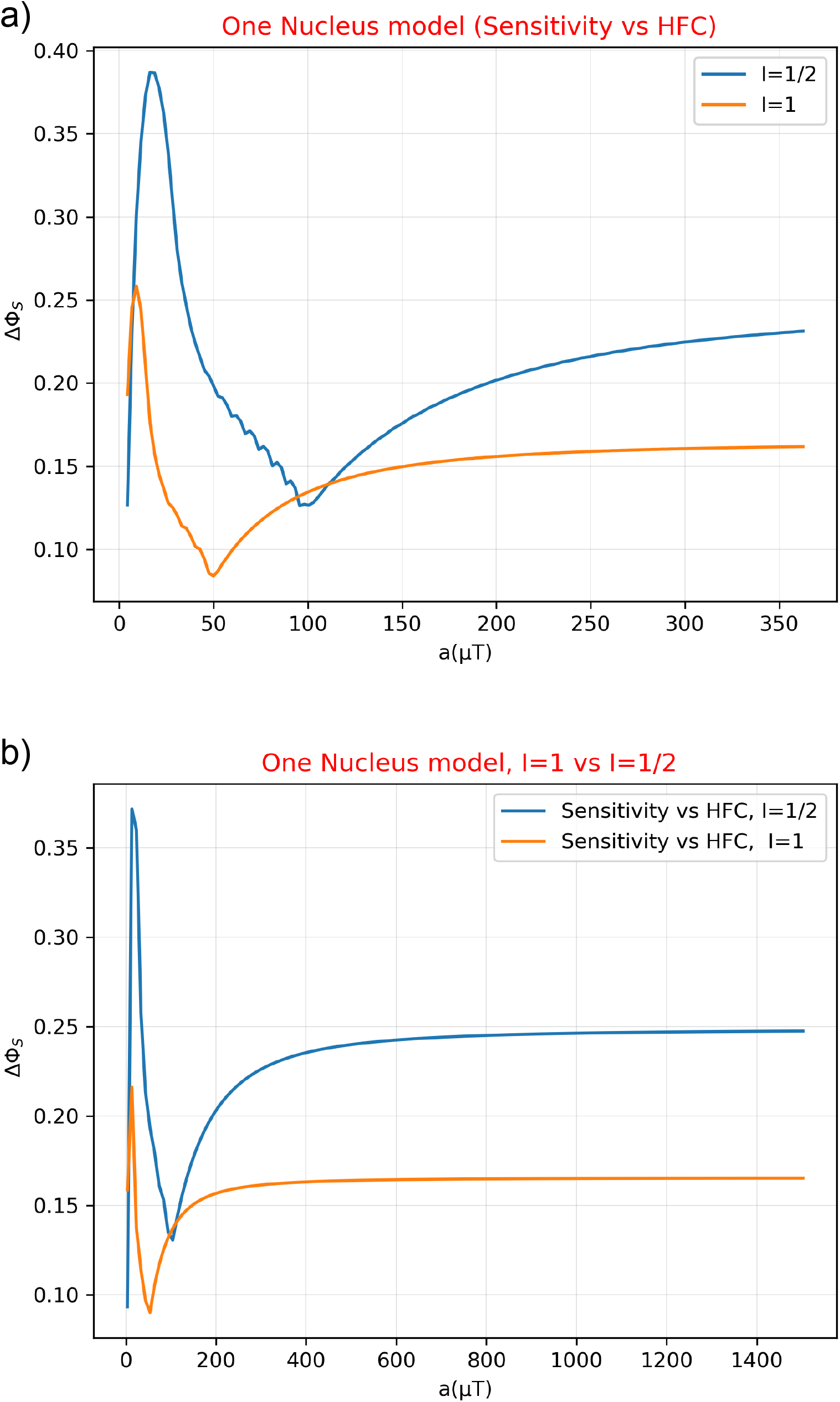
Modelling results for the magnetic sensitivity of cryptochrome (∆Φ_S_) for different spin numbers along a range of anisotropic hyperfine coupling strengths (a). a) Here, the only difference between the treatment of nuclei in the RP model lies in their spin quantum numbers, meaning that we investigate MS with respect to the hyperfine coupling of a single nucleus model for two different spin quantum numbers (*I*=1 and *I*=1/2). In this simple case, we do not include noise in the model. The results demonstrate clear differences between the two spin numbers, with *I*=1/2 giving the greatest magnetic sensitivity for the greatest range of hyperfine coupling constants and the maximum sensitivity for also *I*=1/2 is considerably greater than *I*=1. b) For hyperfine up to 1.5 mT, the sensitivity reaches a plateau, which is equal to 0.16 for *I*=1 and 0.25 for *I*=1/2.

In Figure 4 we again investigate MS for the two different spin numbers across a range of hyperfine coupling strengths, but in this instance we have included noise in the model. As previously specified, our aim was not to investigate the effects of different noise models. Given the calculations by Cai et al. [52] it appears that a specific noise model, such as a perfectly correlated dephasing model, may increase the performance of a chemical compass. We were interested in the effects that this noise model would have on radical pair dynamics for different isotopes. With noise incorporated into the model according to Eq. (4) with Γ= *k* − 3*k* for one nucleus with *I*=1/2, the maximum MS increases from 0.388 (corresponding to a hyperfine coupling strength of 17.4 μT) to 0.411 (corresponding to a hyperfine coupling strength of 20.9 μT). For one nucleus with *I*=1, the maximum MS increases from 0.259 (corresponding to a hyperfine coupling strength of 8.7 μT) to 0.274 (corresponding to a hyperfine coupling strength of 10.6 μT).

**Figure 4.**
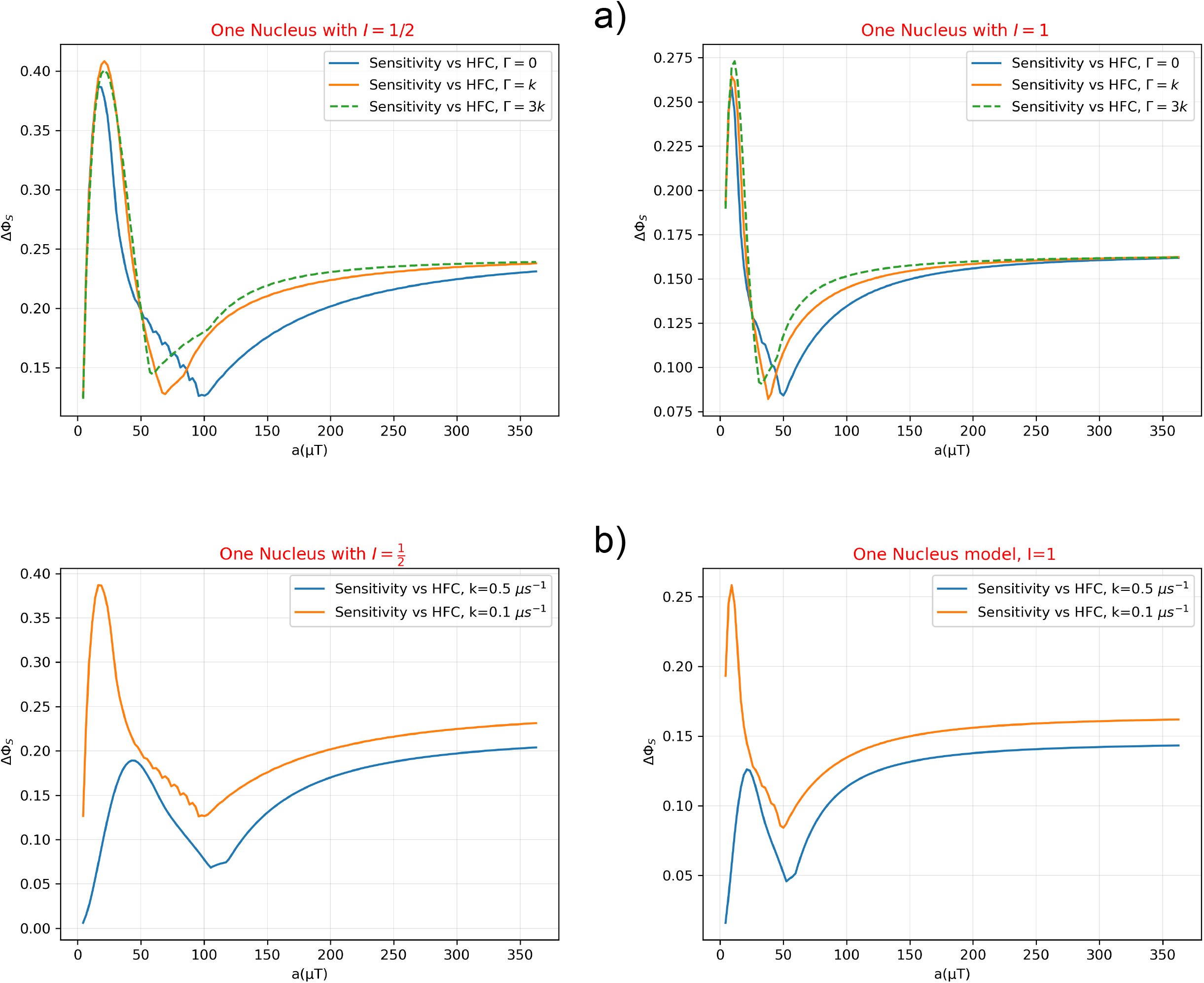
Modelling results for the magnetic sensitivity of cryptochrome (∆Φ_S_) for different spin numbers along a range of anisotropic hyperfine coupling strength (a). a) With noise incorporated into the model, it is shown that MS can be improved for certain hyperfine coupling strengths. For a nucleus with *I*=1/2, the maximum MS increases from 0.388 (corresponding to a hyperfine coupling strength of 17.4 μT) to 0.411 (corresponding to a hyperfine coupling strength of 20.9 μT). For a nucleus with *I*=1, the maximum MS increases from 0.259 (corresponding to a hyperfine coupling strength of 8.7 μT) to 0.274 (corresponding to a hyperfine coupling strength of 10.6 μT). b) It is clear from the results that the sensitivity of the avian compass relies on the recombination rate. A good sensitivity requires a recombination rate that gives enough time for magnetic field effects to occur. Here it is shown that sensitivity decreases when increasing the recombination rate, i.e. shorter lifetime for the radical pair.

### 3.2. Scenario 2

For this second scenario we used the data provided by Jain et al. [43], summarized in Table 2. These authors used three nuclei for each RP of cryptochrome (three nuclei for FAD^•−^ and three nuclei for TrpH^•**+**^) instead of 11 nuclei for each RP as usually considered when investigating hyperfine coupling. For our purposes we focus specifically on the hydrogens H6 and H1 as well as the nitrogens N10 and N1 to compare results for isotopic substitution, see Table 2 for details.

**Table 2.**
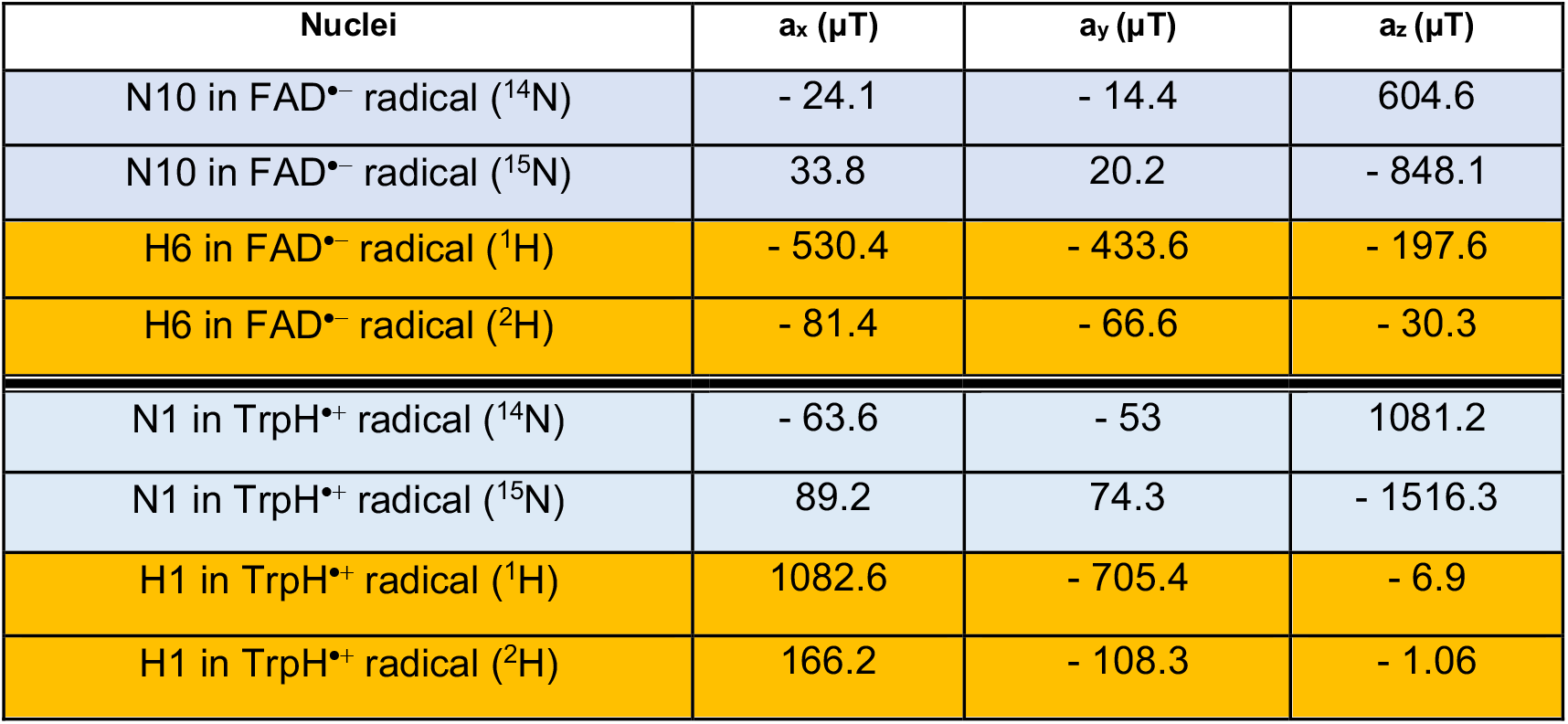
Hyperfine coupling tensors for the [FAD^•−^ - TrpH^+^] radical system, taken from Hiscock et al. (2016). Following Jain et al. (2021), all hyperfine tensors are simultaneously diagonalized.

#### 3.2.1. Scenario 2a

We investigated singlet yield with respect to the inclination (angle describing the orientation of magnetic field to the basis of the hyperfine tensor) of different N and H isotopes in the RP [FAD^•−^ - TrpH^•**+**^] of the cryptochrome molecule, and explored the effect of isotope substitution. The results of these models are shown in Figure 5. It is clear that isotopes with integer spin have consistently higher singlet yield than half integer, even when taking into account their different coupling constants.

**Figure 5.**
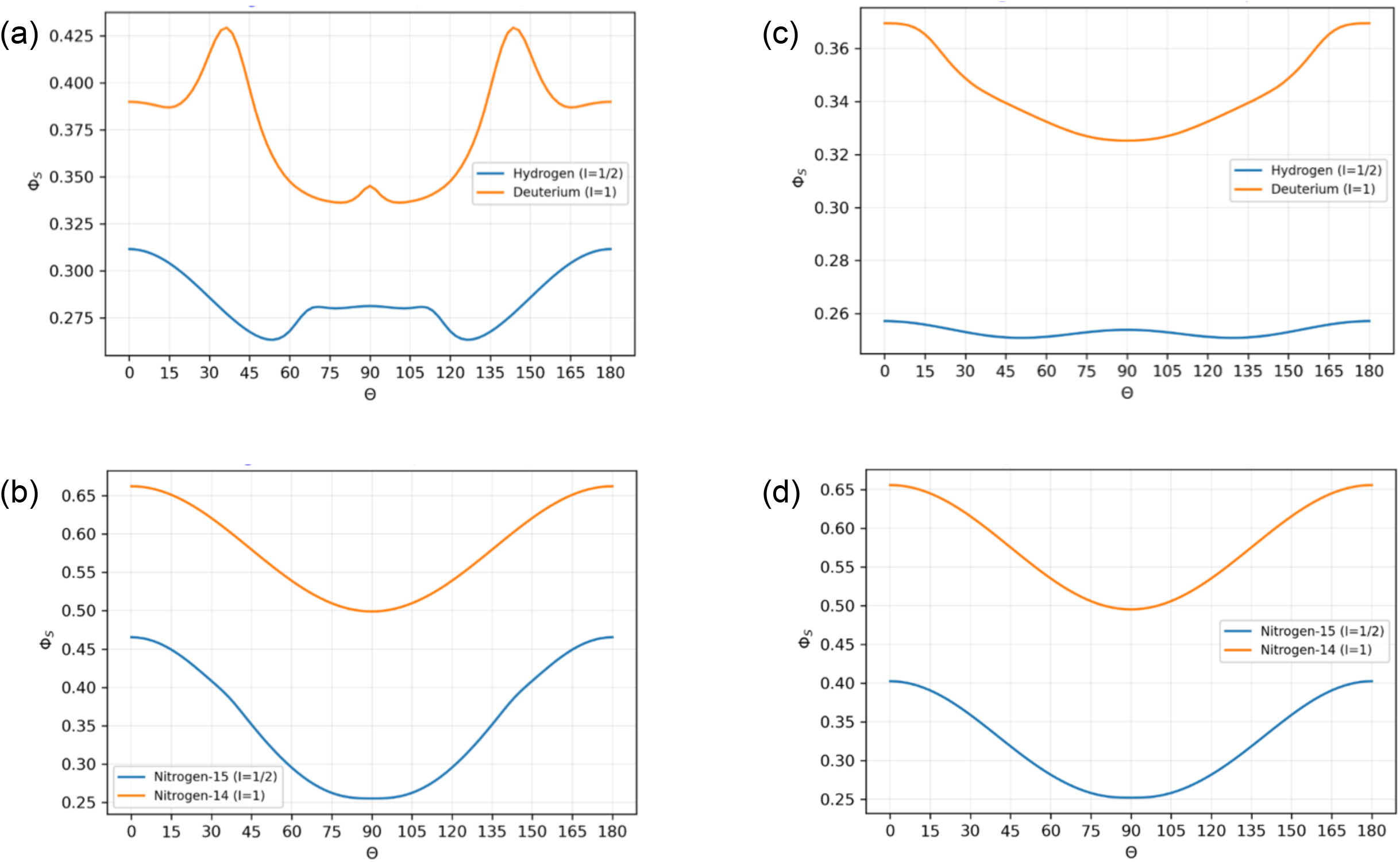
Isotope substitution effects on the magnetic effect (singlet yield, Φ_S_) with different inclinations of cryptochrome for a one-nucleus model. The results are separately shown for the radicals of the RP system in cryptochrome (left panels: FAD^•−^, right panels: TrpH^•**+**^) and for different hydrogen (a: H6, c: H1) and nitrogen (b: N10, d: N1) nuclei and their isotopes. It is clear that isotopes with integer spin have consistently higher singlet yield than half integer, even with different coupling constants. Singlet yield is an interesting feature to consider with respect to RPs in the biological context. In addition to magnetoreception, singlet yield has been used to investigate a number of other biological functions which may depend on radical reactions [62]. In reactions involving reactive oxygen species, singlet yield might be used as an indication of oxidative stress. The strong isotope dependence of singlet yield demonstrated in our results is potentially interesting with respect to observations made that isotopic changes in diet lead to oxidative stress [9].

Similar conclusions are obtained when investigating a two-nuclei model instead of single nuclear spin coupled to one of the electrons (Figure 6). In the two-nuclei model, there are four different combinations of isotopes: ^1^H^14^N, ^1^H^15^N, ^2^H^14^N and ^1^H^15^N. The Hamiltonian now gains an additional hyperfine term:

**Figure 6.**
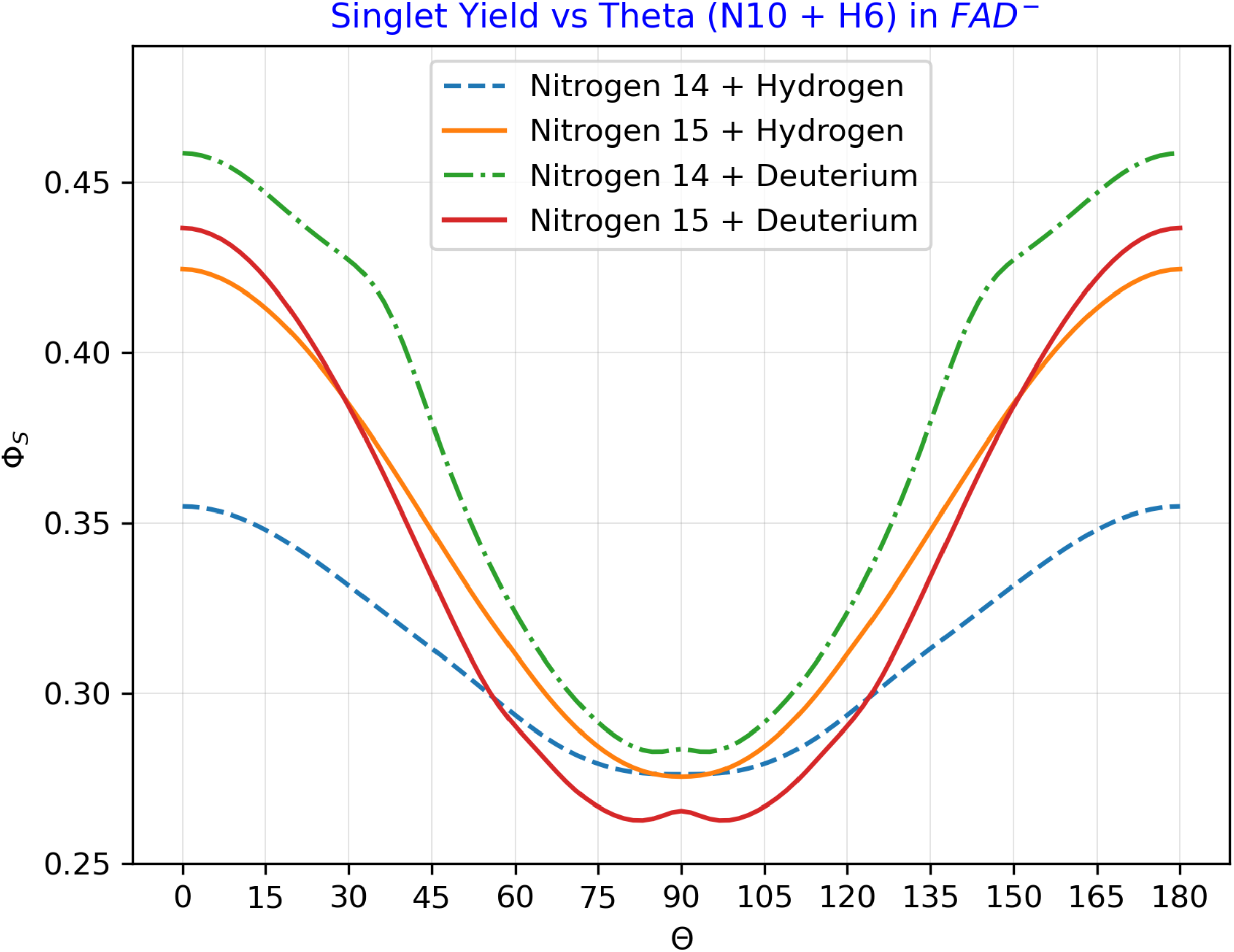
Isotope substitution effects on the magnetic effect (singlet yield, Φ_S_) with different inclinations of cryptochrome for two-nuclei model. The results for the four different combinations of hydrogen and nitrogen isotopes for the two-nuclei model are shown. Results are relatively similar to those of the single-nucleus model (Figure 5), with a minimum at about 90º singlet yield. Singlet yield is greatest for the case in which both nuclei have integer spin, that is for N14 and deuterium.

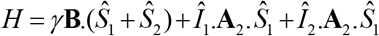

Singlet yield is an interesting feature to consider with respect to RPs in the biological context. In addition to magnetoreception, singlet yield has been used to investigate a number of other biological functions which may depend on radical reactions. In reactions involving reactive oxygen species, singlet yield might be used as an indication of oxidative stress. The strong isotope dependence of singlet yield demonstrated in our results is potentially interesting with respect to observations that isotopic changes in diet lead to oxidative stress [9].

#### 3.2.2. Scenario 2b

Here we investigated MS with respect to the strength of Earth’s magnetic field of different N and H isotopes in the RP [FAD^•−^ - TrpH^•**+**^] and explored the effect of isotope substitution. Results are shown in Figure 7. Scenario 1 (see Figure 3) showed a clear difference between how spin number changes MS, with *I*=1/2 conferring greater MS than *I*=1. However, the effect of the spin number is also dependent on the hyperfine coupling strength, with *I*=1/2 performing best at larger coupling constants than *I*=1. In Scenario 2 it is more difficult to conclude that there is a favourable spin number with respect to MS. This is likely due to the influence of the specific hyperfine coupling constants. While the hyperfine coupling strength decreases with isotope substitution in hydrogen, it increases with isotope substitution in nitrogen. If we consider only the magnetic field strength relevant to migration, which is the geomagnetic field (25-65 µT), then for the case of both hydrogens, *I*=1 gives greater magnetic sensitivity than *I*=1/2. The case is less clear for nitrogen where both *I*=1/2 and *I*=1 show greater MS for specific magnetic field strengths over the geomagnetic window (25-65 µT). This could be due to the fact that although the coupling strengths for the nitrogen isotopes are further from the ideal values, the isotopic substitution from *I*=1 to *I*=1/2 confers some advantage.

**Figure 7.**
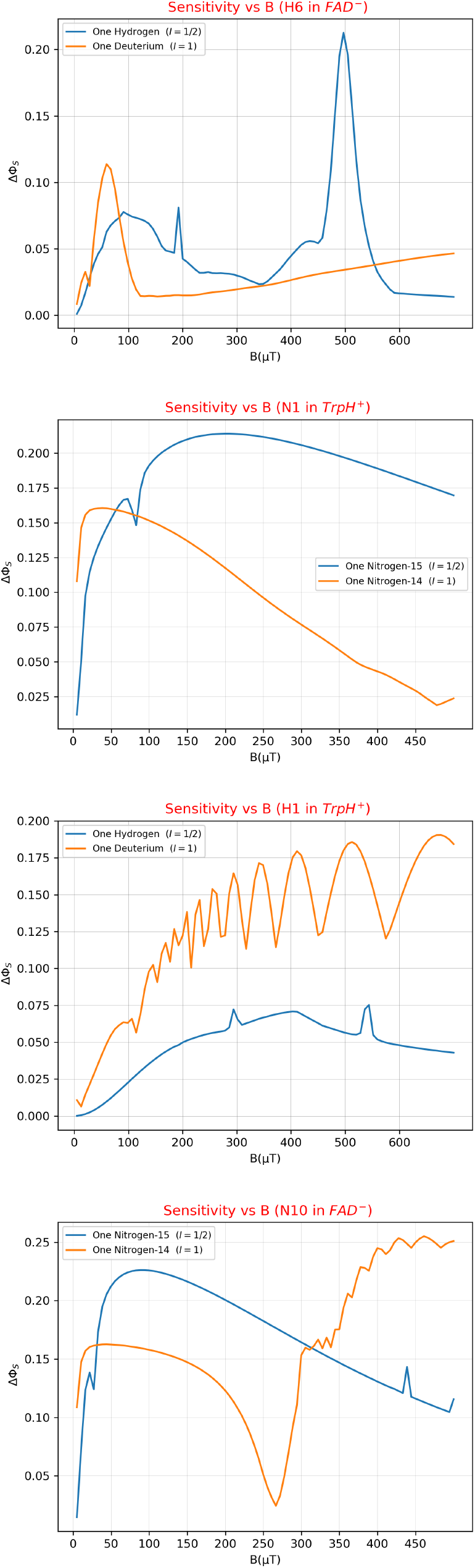
Isotope substitution effects for the magnetic sensitivity of cryptochrome (∆Φ_S_) along a range of strength values of an external magnetic field (B). Examples of different nuclei in the radicals of the RP system in cryptochrome are shown (from to down panels: H6 in FAD^•−^, N1 in TrpH^•**+**^, H1 in TrpH^•**+**^, N10 in FAD^•−^). **Scenario 1** (see Figure 3) showed a clear difference between how spin number changes MS, with *I*=1/2 conferring greater MS than *I*=1. However, the effect of the spin number is also dependent on the hyperfine coupling strength, with *I*=1/2 performing best at larger coupling constants than *I*=1. In **Scenario 2** it is more difficult to conclude that there is a favourable spin number with respect to MS. This is likely due to the influence of the specific hyperfine coupling constants. While the hyperfine coupling strength decreases with isotope substitution in hydrogen, it increases with isotope substitution in nitrogen. If we consider only the magnetic field strength relevant to migration, which is the geomagnetic field (25-65 µT), then for the case of both hydrogens, *I*=1 gives greater magnetic sensitivity than *I*=1/2. The case is less clear for nitrogen where both *I*=1/2 and *I*=1 show greater MS for specific magnetic field strengths over the geomagnetic window (25-65 µT). This could be due to the fact that although the coupling strengths for the nitrogen isotopes are further from the ideal values, the isotopic substitution from *I*=1 to *I*=1/2 confers some advantage.

## 4. Discussion

Before any definite conclusions can be made, it should be acknowledged that the model we have used is a toy model and is the simplest possible interaction of the RP. It serves, however, to illustrate the hypothesis that isotopes may play a distinct and definite role in macroscopic biological outcomes. This influence is exerted in various ways. Our theoretical results show that the spin quantum number (*I*) of a nucleus in an RP strongly influences the dependency of its MS on the strength of the hyperfine coupling. Considering a wide range of hyperfine coupling values from 0 to 350 µT, following Lee et al. [55], the maximum MS is considerably higher, and the asymptote is reached at larger hyperfine coupling values, when *I*=1/2 (∼20 µT) as compared to *I*=1 (∼10 µT). These ideal hyperfine coupling strengths are considerably smaller than those given for the specific nuclei of the RP mechanism, see Table 2. Whether this distinction between *I*=1/2 and *I*=1 can be generalized to non-integer and integer spin would depend on the inclusion of higher spin isotopes. RP performance further improves when the noise model is perfectly correlated, suggesting that RPs in cryptochrome, as well as other biological processes, might be optimized by the unavoidable noise that is present in the environment, in accordance to Bandyopadhyay et al. [57].

Our results for the specific RP [FAD^•−^ - TrpH^•**+**^] of cryptochrome also show strong differences in the dependency of the magnetic field effect (i.e., singlet yield) on inclination, between isotopes with different spin numbers (^1^H vs ^2^H and ^15^N vs ^14^N). In both FAD^•−^ and TrpH^•**+**^ radicals, the H and N isotope with *I*=1 (i.e., ^2^H and ^14^N) consistently experience higher singlet yield along the whole inclination range than their counterparts with *I*=1/2.

The marked isotope-dependence of singlet yield might also offer a novel explanation for isotope-induced oxidative stress, as reported in Zubarev [9]. The role that radical reaction intermediates might play in the chemistry and biology of reactive oxygen species is a growing field of research. Reactive oxygen species (ROS) include a wide variety of oxidant molecules with different properties and biological functions that range from signalling to oxidative damage [58]. In this context, the RP mechanism might give some insight. Usselman et al. show how yields of ROS in live cells are changed by RP dynamics, in particular coherent singlet-triplet mixing [59,60]. For an RP involving oxygen, the substitution of isotopes, which we have here demonstrated to change singlet yield, could thus change the balance of ROS, potentially causing oxidative stress. This is of direct relevance to the migration evolution hypothesis posed here, because migration is a physiologically costly strategy and, consequently, species of birds that conduct long-distance migration have comparatively higher levels of antioxidant resources [61]. It is likely, then, that the possible role of isotopes on the evolutionary origin of migration also leads to differences in oxidative stress between migratory and non-migratory animals, something that should be explored in the future. The role that isotopes play in RP-mediated changes to singlet yield have also been investigated in a number of other biological contexts, including consciousness [62]. It would thus seem important to determine how isotopes facilitate or inhibit the development of biological functions that may confer some evolutionary advantage on an organism.

It is interesting to note, however, that the greater singlet yield of nuclei with *I*=1 does not necessarily translate into greater magnetic sensitivity, as demonstrated by the results in Figure 7, for the MS of the H and N isotopes investigated in FAD^•−^ and TrpH^•**+**^. We consider in particular the range of external magnetic field values corresponding to the geomagnetic field (25-65 µT). For both hydrogens, in contrast to the results given by the simple case investigated in Scenario 1 (Figure 3), MS is improved by isotope substitution with *I*=1. This is likely due to the hyperfine coupling environment. From the results in Figure 3 it appears that nuclei with *I*=1 perform better at smaller coupling constants. Isotopic substitution in hydrogen gives smaller coupling constants. However, further investigation would have to be done before definite conclusions are drawn. The case for nitrogen and its isotopes is less clear. Over the relevant geomagnetic window (25-65 µT), both *I*=1/2 and *I*=1 are more favourable at certain values of the magnetic field. For the specific nitrogen N10, the isotopic substitution with *I*=1/2 dramatically improves the MS at certain values of the field. While it is difficult to draw any absolute conclusions, it is potentially interesting that for both hydrogen and (at least partly) nitrogen, isotopic substitution acts to increase the MS. Although we have investigated here only isotopes of H and N, differences in the potential to contribute to the performance of cryptochrome as a magnetic compass are likely to exist also between isotopes of the other constituent elements of the molecule (i.e., C and O), as well as any element functioning in other biochemical processes (e.g., S and P). The fact that heavy isotopes are less abundant in nature than their lighter counterparts (in the case of δ^2^H and δ^15^N this being only 0.0115% and 0.364%, respectively, see Table 1) does not preclude the possibility that differences in the isotopic composition of a biomolecule such as cryptochrome between species of animals generate differences in the performance of the involved biochemical process, despite the negligible effect of heavy isotopes on the speed of chemical reactions that is predicted by conventional chemical kinetics. The Isotopic Resonance Hypothesis, on the other hand, posits that optimal, ‘resonance’ abundances of stable isotopes reduce the complexity of biochemical systems, affecting the kinetics of direct and reversed reactions within the system and maximizing the efficiency of biochemical reactions [13,14,63]. The Isotopic Resonance Hypothesis has received empirical support from enzymatic reactions, also mediated by a quantum process such as quantum tunnelling [64,65]. Thus, the enzymatic catalytic action of luciferase on its substrate luciferin with a deuterium concentration in the local environment 2-4 times higher than the normal concentration of 150 ppm but still considerably low (250-350 ppm) exhibits a significant change that is not predicted by conventional chemical kinetics. This indicates that small concentrations of heavy isotopes exert disproportionately strong effects on the kinetics of enzymatic processes [66]. Similarly, animals with a cryptochrome composition enriched in isotopes of, for example, hydrogen and nitrogen may have a superior magnetic sensitivity, this not being hindered when such isotopes are heavy, such as deuterium (^2^H).

Consequently, our results show that the isotopic composition of cryptochrome can exert significant effects on the performance of RPs, hence influencing the likelihood of developing an ability for magnetoreception and, thus, for migration. This means that the evolution of migration may be favoured only under certain isotopic compositions of cryptochrome. This has never been proposed before. But, indeed, non-adaptive origins of adaptive traits are relevant sources of evolutionary innovations, particularly for metabolic traits [67].

Physiology is considered to have a foundational role in migration, which has been studied in detail (e.g., [68]). The evolutionary maintenance of migration is based on its adaptive benefits, as it allows animals to get trophic resources that would otherwise be inaccessible, and is also well understood. However, it is unknown whether differences in migration among species are due to genetic or environmental effects [69,70]. Recently, a study on a single migratory bird species (the peregrine falcon *Falco peregrinus*) has revealed that divergence in the gene *ADCY8* explains differences in migration distance between populations, but not why these birds always migrate in some extent [71]. As a consequence, the ultimate factors that generally drive animal migration and that explain the evolutionary appearance of this behaviour in some species and not in others remain undeciphered [72]. Interspecific differences in the isotopic composition of cryptochrome may therefore represent a general explanation for the origin of animal migration.

An effective test of our hypothesis will require empirical comparisons of cryptochrome isotopologues between migratory and non-migratory species of birds or other animals. Some already existing empirical results, however, may support our proposal. In fact, an investigation on the isotopic composition of hair keratin in three species of coexisting bats resulted in an isospace (δ^13^C vs. δ^15^N) of a long-distance migratory bat (*Pipistrellus nathusii*) markedly distinct from the isospaces of non-migratory and mid-distance migratory bats (*Pipistrellus pipistrellus* and *Nyctalus noctula*) [73]. Considering that the isotopic profile may be consistent among all molecules of a species of animal (see Introduction), the latter results may be in accordance with the hypothesis proposed here.

A causal association between animal isotopic profiles and their macroscopic characteristics would represent a non-adaptive and non-organic source of evolutionary innovations, and a connection between the mineral and the organic world, unveiling attributes of organisms not carried by their genetic code. The transference of atomic information across organisms would represent a DNA-independent inheritance that has never been investigated, but future studies should investigate if this may in fact be the evolutionary origin of animal migration and other biological processes.

## Acknowledgements

We would like to thank Ilya Sinayskiy for his helpful discussions and comments during this work.

## Ethics

No ethical issues are associated with this study.

## Data, accessibility

This article has no additional data.

## Authors’ contributions

I.G. conceptualised the role of isotopes in organic evolution. A.H., B.A. and F.P. contributed to physics interpretation and model development. A.H. did the calculations and spin dynamics simulations. I.G., A.H., B.A. and F.P. wrote and reviewed the manuscript.

